# Optimal Maturation Protocols for High-Affinity Antibody Targets: A Path-Integral Approach

**DOI:** 10.64898/2025.12.21.695799

**Authors:** Marian Huot, Marco Molari, Rémi Monasson, Simona Cocco

## Abstract

Affinity maturation is a stochastic evolutionary process allowing the adaptive immune system to produce B-cells capable of recognizing antigenic molecules. One of the main factors influencing the quality of the maturation outcome, quantified by the affinity of the produced antibodies to the antigen, is the time-course of antigen availability. In this paper, we introduce a stochastic model for affinity maturation, calibrated against in vivo lineage dynamics and deep mutational scanning data, and address the following question: what is the best antigen-concentration protocol maximizing the probability to reach a target affinity value at the end of the process? We introduce a path-integral formalism to identify the maturation trajectories of the rare clones ending at the desired target affinity, estimate their probabilities, and maximize them over the antigen-concentration temporal profile. The theoretical optimal concentration protocol is approximated by a discrete three-injection schedule; we show that such temporal modulation of selection pressure outperforms constant-dosage regimes.

## I. INTRODUCTION

Vaccination remains one of the most effective interventions in medical history, yet the development of potent immunogens against highly mutable pathogens, such as HIV and influenza, continues to defy conventional empirical approaches. The efficacy of a vaccine relies fundamentally on its ability to trigger Affinity Maturation (AM), a rapid Darwinian evolutionary process occurring within Germinal Centers (GCs). In these microanatomical structures, B cells undergo iterative rounds of somatic hypermutation and selection, progressively evolving B-cell receptors (BCRs) capable of binding antigen with high affinity [1–4]. While the biological mechanisms governing this process—clonal expansion, mutation, and competitive selection for survival signals—are well characterized, the quantitative principles that determine the optimal evolutionary trajectory of a B-cell lineage remain an active area of theoretical investigation.

A central variable governing the efficiency of this evolutionary search is the availability of antigen. Antigen concentration acts as a tunable control parameter for selection pressure [5–7]: it determines the energetic threshold a B cell must overcome to capture antigen, present it to T follicular helper cells, and receive survival signals. Previous computational and experimental studies have highlighted a fundamental trade-off regarding antigen dosage [8–10]. High concentrations relax selection pressure, allowing low-affinity clones to survive and dominate the population, thereby stalling the maturation process. In contrast, strict selection mediated by low antigen concentrations accelerates affinity gains but increases the risk of population extinction before high-affinity variants can emerge [11]. It is therefore important to identify the best antigen administration protocols, maximizing the likelihood of reaching a high-affinity final state.

We address the problem of optimal immunization by leveraging established computational models of affinity maturation [10–14]. Despite their idealized nature, these frameworks effectively encapsulate the stochastic fluctuations and complex selection pressures that define the process. We use these models to ask: how can we quantify—and subsequently maximize—the probability of reaching a desired affinity through the modulation of antigen availability? Specifically, we investigate the following: what is the probability that a B-cell lineage achieves a target affinity, and how can the temporal profile of antigen be tuned to maximize this likelihood? Our objective is to identify the most probable evolutionary trajectory leading from a wild-type state to a specific affinity threshold, along with its associated probability of success.

To do so, we introduce a mathematical framework that treats affinity maturation as a stochastic trajectory optimization problem on a fitness landscape. We map the discrete, agent-based dynamics of B-cell evolution onto a continuous path-integral formalism to identify the “least-action” trajectories—the most probable evolutionary histories leading to high-affinity states. We also take into account the fluctuations around these least-action trajectories. This formalism allows us to analytically derive the time-dependent antigen concentration that maximizes the probability of evolving high-affinity clones while respecting biological constraints such as population viability and carrying capacity.

We calibrate our model using time-resolved lineage data and deep mutational scanning profiles from in vivo experiments, ensuring that our theoretical predictions are grounded in realistic biophysical parameters. Contrary to the common intuition that stronger selection always yields better antibodies, we show that the optimal protocol requires a dynamic modulation of selection pressure that keeps the population near a critical “tipping point” of extinction. We further demonstrate how this theoretically optimal continuous solution can be discretized into a practical multi-injection vaccination schedule, providing a rational basis for the design of immunization protocols that maximize evolutionary gain.

## II. MODEL AND COMPARISON WITH EXPERIMENTAL DATA

### A. Affinity maturation model

We describe a stochastic model for antibody affinity maturation within a single Germinal Center (GC). The variable component of the B-cell receptor affinity is quantified by the affinity parameter *ϵ*, measured in units of *k*_*B*_*T*, where *k*_*B*_ is the Boltzmann constant and *T* is the organism’s temperature. This choice of units allows one to simply express Boltzmann factors as *e*^*ϵ*^. This affinity is related to the dissociation constant, *K*_*D*_, between the B-cell receptor and the antigen through *ϵ* = −log(*K*_*D*_) + *ϵ*_*a*_ where *ϵ*_*a*_ is the wildtype log dissociation constant. Here, *K*_*D*_ is expressed in nanomolar (nM) units; choosing different units would shift the values of *ϵ* by a constant amount.

To capture site-specific granularity, we define the B-cell affinity as the sum of these individual site contributions:

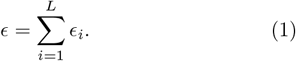

In particular, the wildtype (unmutated) B-cell has *ϵ*_*i*_ = 0 for all sites. The maturation process proceeds through iterative rounds of duplication, mutation, and selection (Figure 1 and [10, 12]) that modify the values of the *ϵ*_*i*_’s.

**FIG. 1.**
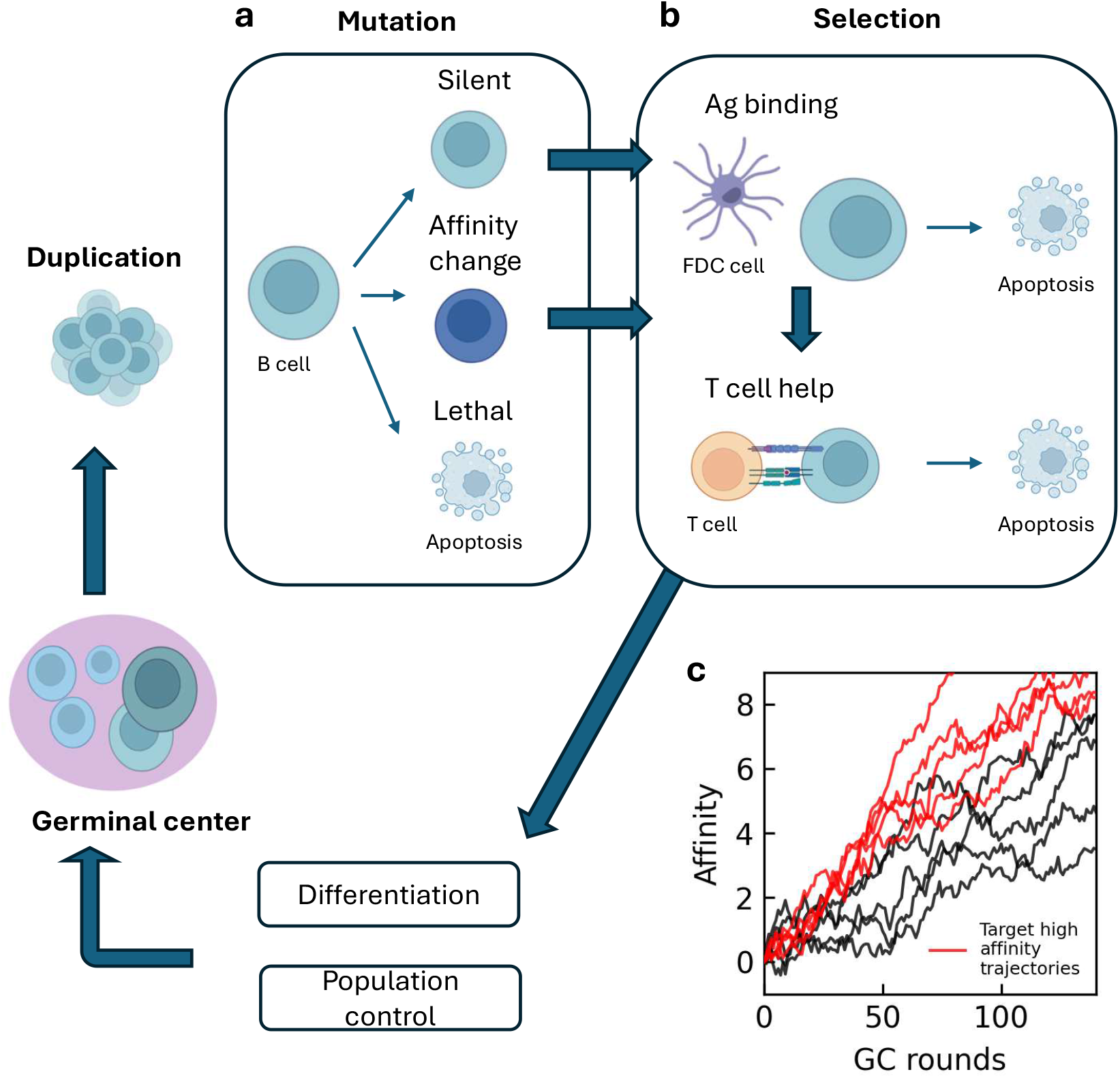
Schematic of the affinity maturation model. The simulation proceeds through iterative rounds of evolution. **a)** Duplication and somatic hypermutation: B cells proliferate and mutate, with outcomes classified as silent, lethal, or affinity-affecting. **b)** Selection and regulation: Cells undergo a two-step selection process involving antigen binding and competition for T cell help. The population is subsequently regulated via differentiation and random removal to enforce carrying capacity. c) Stochastic affinity trajectories of B cells over GC rounds. The red curves highlight the subset of target trajectories that successfully reach high-affinity states, which are the focus of this study.

Cells first duplicate in the GC Dark Zone and then mutate one site with probability *µ* for each sequence (Figure 1a). Mutation effects can be silent (with probability *q*_*sil*_), lethal (*q*_*let*_, resulting in the immediate removal of the cell), or affinity affecting (*q*_*aa*_) [10, 12, 14] (see Table I). In the latter case, we update a random site *i* by a random additive contribution 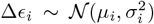. We make these contributions site-specific, with the intention of setting them from experimental measurements.

**TABLE 1.** Parameter values used for the simulations.

| Symbol | Reference | Description |
| --- | --- | --- |
| $L = 40$ | [15] | Number of CDR residues |
| $T = 140$ | [15] | Number of GC rounds |
| $\epsilon_a = \log(40)$ | [15] | $\log K_d(\text{WT})$ (nM) |
| $\Delta\epsilon_i \sim \mathcal{N}(\mu_i, \sigma_i^2)$ | [15] | Effect of site mutations |
| $\mu = 0.001 \times L \times 3$ | [12] | Mutation probability per sequence of size $L$ |
| $q_{\text{sil}} = 0.3$ | [10, 12, 14] | Probability of silent mutation |
| $q_{\text{let}} = 0.5$ | [10, 12, 14] | Probability of lethal mutation |
| $q_{\text{aa}} = 0.2$ | [10, 12, 14] | Probability of affinity-affecting mutation |
| $N_{\text{max}} = 2500$ | [10] | GC max population size |
| $C$ | free parameter | Antigen concentration (nM) |

As a consequence, each daughter cell keeps same affinity with probability *p*_sil_ = (1 − *µ*) + *µ q*_*sil*_, dies with probability *p*_let_ = *µ q*_*let*_, or modifies affinity with probability *p*_aa_ = *µ q*_*aa*_.

Following mutation, cells migrate to the Light Zone to undergo selection (Figure 1b). In the first selection step, B-cells attempt to bind the antigen (Ag) exposed on Follicular Dendritic Cells. The probability of successfully binding and surviving this step depends on the antigen concentration *C*(*t*), expressed in nMol, and the total binding energy. Cells that fail to bind undergo apoptosis. The probability of antigen binding is:

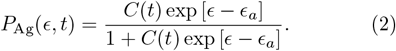

Surviving cells compete for survival signals provided by T-cells. This introduces a competitive element where B-cells with higher affinity are preferentially selected. The probability of surviving this selection step is:

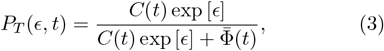

where 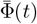 represents the mean fitness of the current population, acting as a competitive threshold. It is defined as the population average of the affinity term:

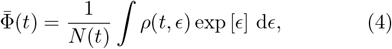

where the cell density *ρ*(*t, ϵ*) represents the expected number of cells with affinity *ϵ* (per unit of affinity) at time *t*. The total population size is given by the integral

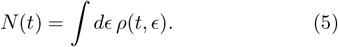

The 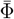-dependent term at the denominator of Eq.(4) ensures that the selection pressure adapts dynamically to the improving affinity of the population.

The surviving cells differentiate with uniform probability *p*_diff_ = 10%. Cells that differentiate are removed from the population.

Finally, if the population exceeds the maximum carrying capacity *N*_max_, cells in excess are randomly removed, irrespective of their affinity [10]. Iterating these GC cycles yields stochastic trajectories of affinity maturation, as illustrated in Figure 1c.

### B. Comparison with experimental affinity maturation trajectories

We calibrate the model parameters in this stochastic algorithm using experimental data from DeWitt et al. [15], who investigated the reproducibility of evolutionary trajectories by “replaying” germinal center reactions starting from a monoclonal B-cell population. In their study, B cells specific for the chicken IgY antigen were transferred into GC-deficient mice, which were subsequently immunized with IgY to initiate the affinity maturation process. Following immunization, the authors reconstructed the phylogenetic histories and affinity trajectories of B-cells from over one hundred individual GCs. Furthermore, they utilized Deep Mutational Scanning (DMS) to quantify the effects of virtually all possible single-amino acid substitutions on the receptor’s binding affinity.

We directly incorporate these empirical measurements into our model parameters. Specifically, the activation energy *ϵ*_*a*_ is derived from the measured wildtype affinity to antigen, and the stochastic update factor Δ*ϵ*_*i*_ is sampled from a site-specific normal distribution 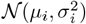 fitted on the amino acid mutational effects determined by their DMS experiments (Table I). We restrict our analysis to mutations that are one nucleotide away from wild-type. We focus our analysis on the *L* = 40 CDR residues as they exhibit significantly higher mutability in both light and heavy chains compared to other residues [15].

Then, we validate our simulation by reproducing the population-level affinity maturation trajectories observed in their time-course sequencing data. We first compare the mean affinity evolution of our stochastic simulation (averaged over 10 independent realizations) against the experimental measurements (Figure 2a). By adjusting a single free parameter—a constant antigen concentration *C*(*t*) = *C*_*exp*_—our model accurately captures the dynamics of affinity gain, reproducing the characteristic rapid initial rise followed by a slower improvement observed *in vivo*.

**FIG. 2.**
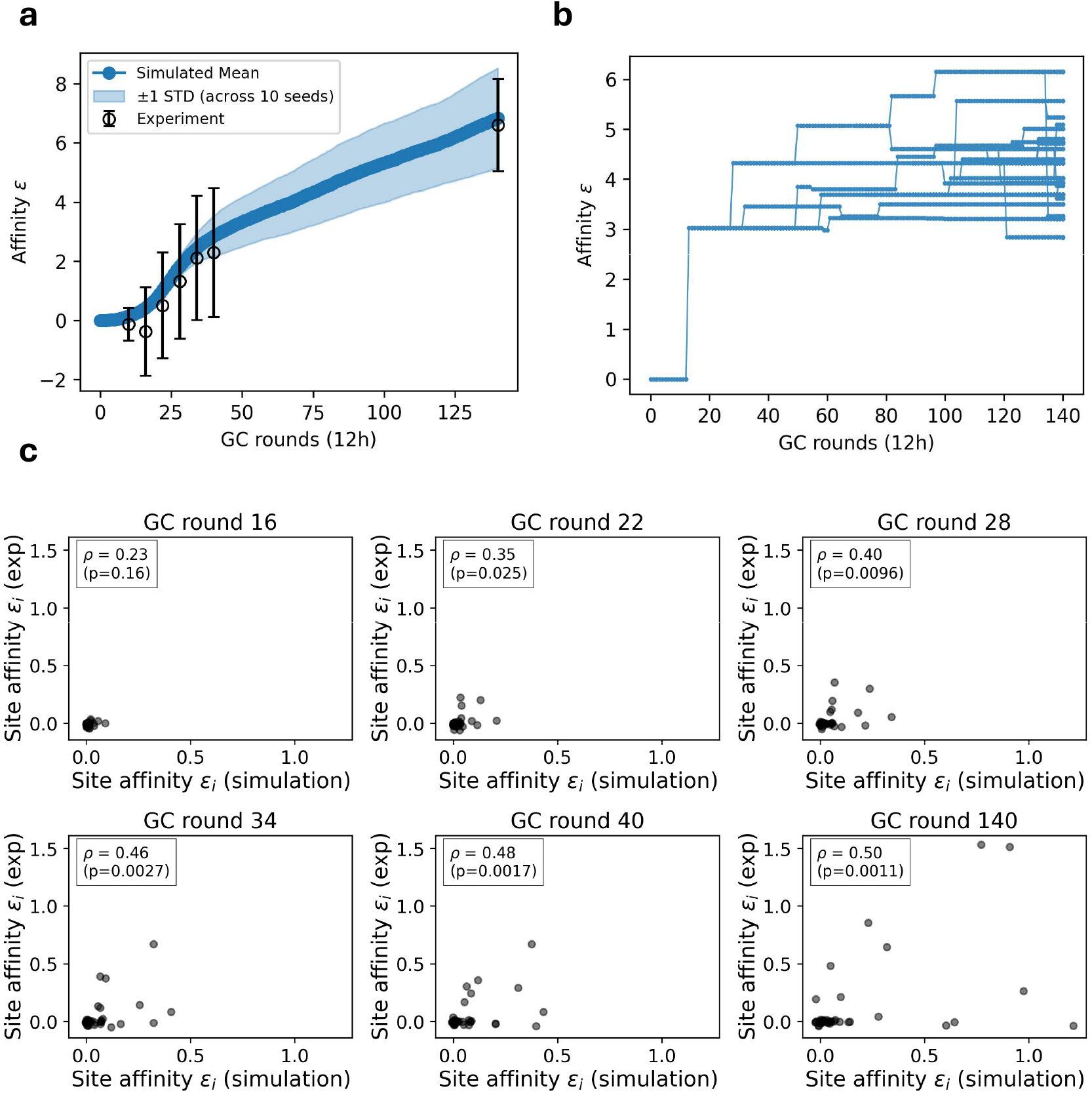
Comparison with experimental data from DeWitt et al. **a)** Comparison of simulated affinity trajectories (mean of 10 independent stochastic realizations) against experimentally measured affinity maturation in DeWitt et al. Only one free parameter (antigen concentration *C*) was adjusted to match the experimental results. **b)** Evolutionary tree of a single founder cell, with daughter cells observed at the final time step. B cells with lower affinities relative to the population average are predominantly observed only at the sampling endpoint. **c)** Comparison of simulated and experimental site-specific affinities at different time points of the maturation process. Pearson correlation coefficient with p value are indicated.

To investigate the lineage dynamics underlying these trajectories, we reconstructed the phylogenetic history of a representative simulated founder cell (Figure 2b). The resulting tree topology reveals a distinct distribution of fitness states: while the backbone of the lineage is dominated by high-affinity clones, B-cells with lower affinities compared to the population mean are predominantly observed only at the final time steps (the leaves of the tree). This observation is consistent with the “pull of the present” effect described by DeWitt et al., where recent deleterious mutations are sampled simply because selection has not yet had time to purge them from the population. Conversely, the historical nodes show a survivorship bias (“push of the past”) where only fitness-increasing mutations tend to fixate and leave descendants.

Furthermore, we demonstrate that our model preserves the statistical properties of site-specific affinity contributions. To do so, we compare the effects of single mutations observed in experiments with the site-specific affinity contributions derived from our simulations (Figure 2c). We observe strong agreement across different time points, with Pearson coefficient reaching *ρ* = 0.5 (p–value = 0.0011) at the end of simulation.

## III. BACKTRACKING AFFINITY MATURATION TRAJECTORIES

### A. Continuous description and Fokker–Planck equation

We now derive a continuous description of the evolutionary dynamics. To make the problem computationally tractable, we reduce the high-dimensional shape space to a single effective binding energy coordinate, denoted hereafter by the scalar *ϵ*. The mutational effects on affinity of this single site are obtained from a normal distribution fitted on all sites mutational effects for single nucleotide away mutations. In the limit of large population sizes and frequent, weak-effect mutations, the evolution of the binding energy distribution *ρ*(*t, ϵ*) can be approximated by a Fokker–Planck equation describing an advection-diffusion process.

We first decompose the deterministic growth and decay dynamics. The baseline growth rate, *λ*, aggregates the contributions of cell duplication, lethal mutations (occurring with probability *p*_let_), and differentiation (probability *p*_diff_) during a single evolution round:

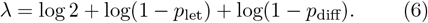

Selection pressures modulate this baseline rate based on the cell’s affinity. The net fitness landscape is shaped by the probability of antigen binding, *P*_Ag_(*ϵ, t*), and the probability of receiving T-cell help, *P*_*T*_ (*ϵ, t*). The total energy-dependent growth rate is therefore:

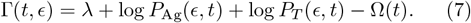

To strictly enforce the carrying capacity *N*_max_, we introduce a pressure term Ω(*t*). This term prevents unbounded growth by penalizing the population growth rate when the global population size *N* (*t*) exceeds the threshold:

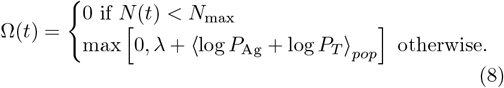

Next, we characterize the stochastic effects of affinity-affecting mutations. While mutations do not alter the instantaneous population size, they redistribute cells along the binding energy axis. In the continuum limit, this jump process is approximated by a drift-diffusion operator defined by drift velocity *v* and diffusion coefficient *D* (see Appendix). Physically, since the majority of random mutations are deleterious, we have a negative drift (*v <* 0) that acts as an entropic force opposing selection.

Combining these contributions, the time evolution of the density *ρ*(*t, ϵ*) is governed by the following Fokker– Planck equation:

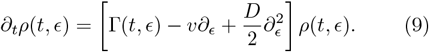

To systematically investigate the impact of antigen availability on evolutionary outcomes, we numerically solved the Fokker-Planck equation over 140 GC cycles under varying antigen concentrations (Figures 3 a-d). The dynamics reveal a critical dependency on the strength of selection pressure. At insufficient antigen concentrations, the stringent selection barrier exceeds the population’s adaptive capacity, leading to rapid extinction (Figure 3a). Conversely, excessively high concentrations create a permissive environment where selection is weak (Figure 3d) [10]. Under these conditions, the population maintains high diversity (large standard deviation) but fails to achieve significant affinity improvement relative to the wildtype. The optimal evolutionary outcome is achieved at intermediate concentrations (Figure 3b and 3c); in this regime, selection pressure is sufficiently strong to drive the population toward high-affinity states, yet permissive enough to sustain population viability, resulting in the most robust maturation. The existence of these three regimes is consistent with the stationary equilibrium of the population dynamics equations for the B-cell maturation process [10].

**FIG. 3.**
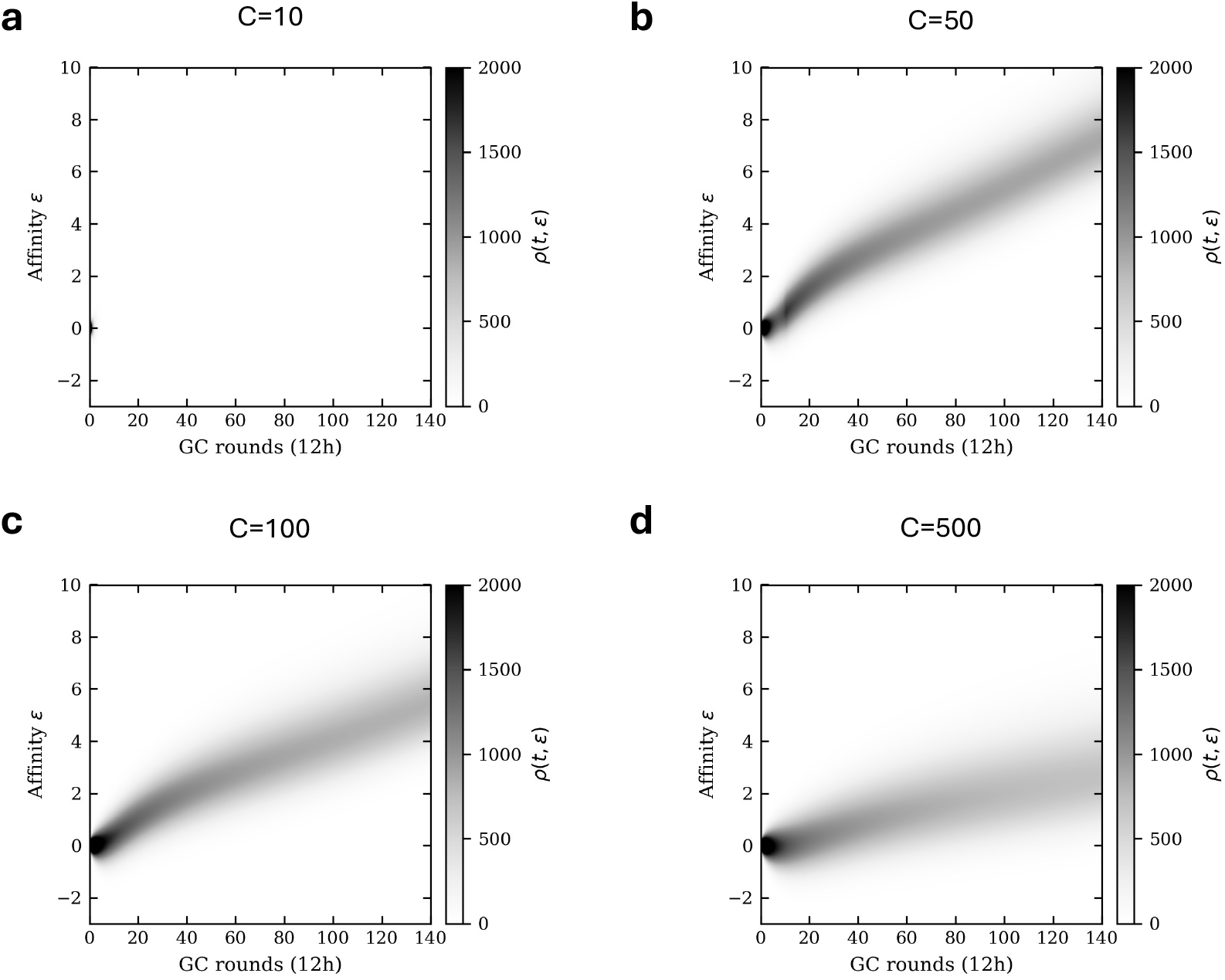
Impact of antigen concentration on affinity maturation outcomes. **a-d)** Temporal evolution of the B-cell affinity density, obtained by numerically solving the Fokker-Planck equation over 140 GC cycles under varying antigen concentrations. Color indicates the probability density at different time steps. Insufficient antigen concentration leads to population extinction, whereas excessively high concentration results in relaxed selection pressure and a lower final affinity of the B-cell population.

### B. Path integral representation and backward trajectories

We next try to provide the shape of the most likely trajectory leading to any final energy state. The infinitesimal generator of the dynamics defined by the Fokker-Planck equation allows us to construct the path integral for the trajectory *ϵ*(*t*) ending at the affinity *ϵ*_*f*_ at the final time *t*_*f*_. After introducing the auxiliary conjugate field to enforce local constraints and integrating it out (see Appendix), we obtain the path-integral representation (Figure 4a):

**FIG. 4.**
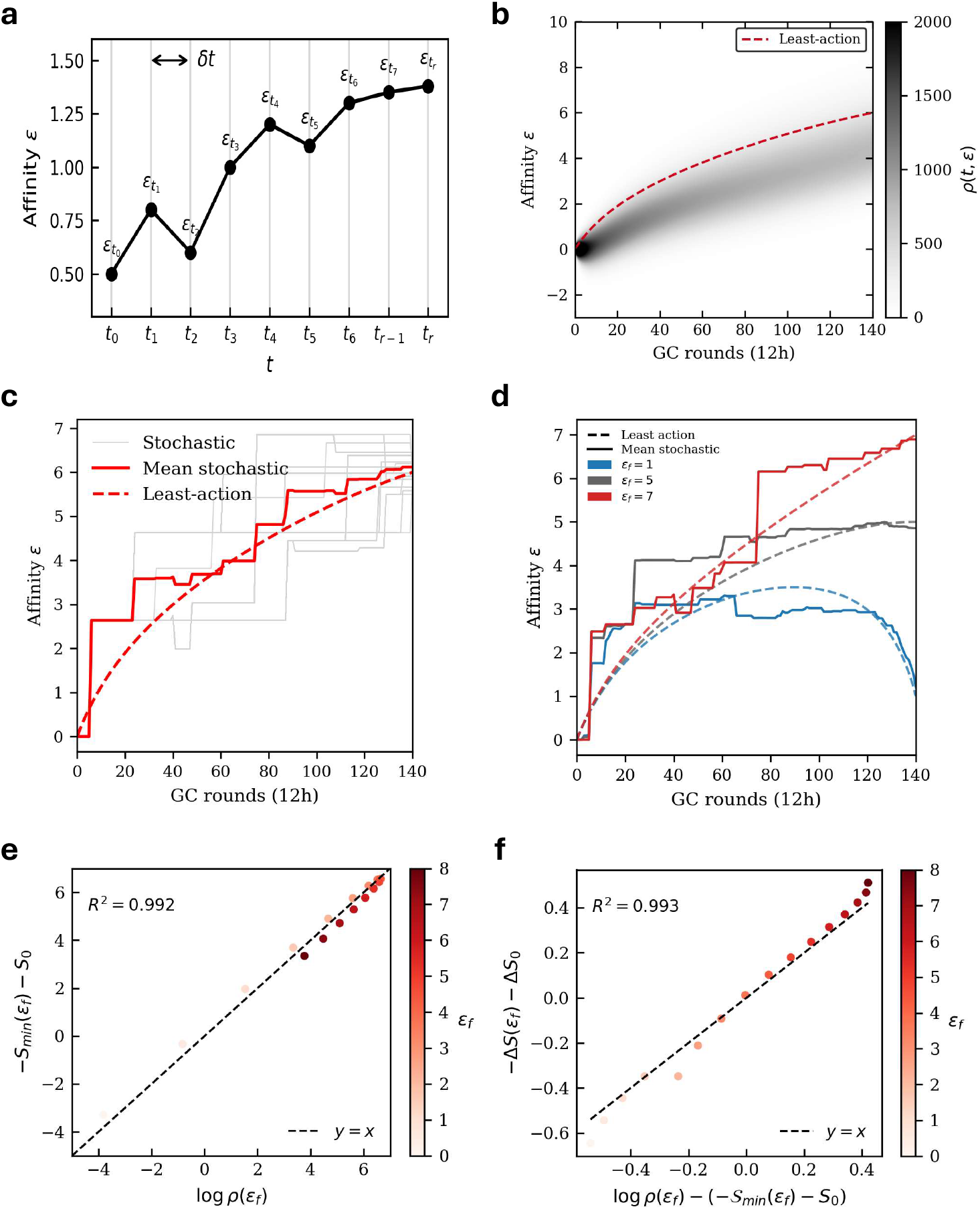
Least-action analysis of affinity maturation trajectories. **a)** Schematic representation of the path integral formulation, where a continuous trajectory is discretized into steps of size *δt*. **b)** Reconstruction of the most likely evolutionary history. The dashed line represents the theoretical least-action trajectory, computed by minimizing the action *S*[*ϵ*] conditioned on a fixed final affinity of *ϵ*_*f*_ = 6. **c)** Comparison of theoretical least-action path and stochastic lineages that successfully evolved to the target affinity *ϵ*_*f*_ ≈ 6. **d)** Comparison of theoretical least-action path and mean of stochastic lineages that successfully evolved to the target affinity *ϵ*_*f*_. **e)** Validation of the least action approximation. The minimum action *S*_*min*_ corresponding to a specific final state scales linearly with the logarithm of the final population density *ρ*(*ϵ*_*f*_), confirming the approximation *ρ* ∝ exp(−*S*_*min*_). *S*_0_ is a constant fixed to the average difference between *S*_*min*_ and *ρ*. **f)** Comparison of the entropic correction −Δ*S* and the error made by the least action approximation, compared to the Fokker Planck density. Δ*S*_0_ is a constant offset introduced to ensure both quantities are plotted on a comparable range.

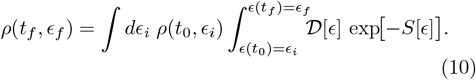

Here, the effective action *S* of a trajectory ***ϵ*** = {*ϵ*(*t*), 0 ≤ *t* ≤ *t*_*f*_} is given by

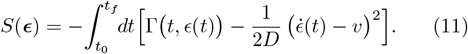

The path integral is dominated by the saddle-point trajectory that minimizes this action. For a population starting with a given affinity *ϵ*_*i*_, the probability density at the final time is approximated by the contribution of this single “most likely” evolutionary path:

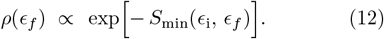

We identify the effective time-dependent Lagrangian,

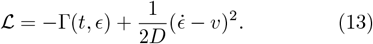

Minimizing the action via the Euler-Lagrange equation,

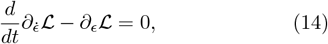

yields the equation of motion for the optimal trajectory:

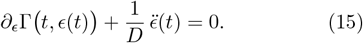

This second-order ODE describes a “particle” moving through the fitness landscape Γ under the influence of an effective inertial term proportional to the inverse diffusion coefficient.

By imposing boundary conditions fixed at the wildtype state *ϵ*(*t*_0_) = 0 and a target *ϵ*(*t*_*f*_) = *ϵ*_*f*_, we numerically solve the corresponding Euler-Lagrange equation (Figure 4b). The resulting least-action trajectory accurately traces the ridge of the probability density obtained from the direct integration of the Fokker-Planck equation, identifying the most probable route through the dynamic fitness landscape. Moreover, we test the predictive power of this theoretical framework against the microscopic dynamics of the discrete model. We isolate the subset of stochastic realizations that successfully reach the target affinity *ϵ* by the final time point (Figures 4c and 4d). Notably, we observe in both the stochastic simulations and the least-action paths that the divergence between high- and low-affinity trajectories becomes particularly pronounced after one month of maturation (Figure 4d). While individual lineages fluctuate due to the stochastic nature of mutation and selection, their ensemble average is remarkably well described by the deterministic least-action solution. This agreement demonstrates that the Euler-Lagrange equation captures the mean evolutionary dynamics of successful lineages.

### C. Probabilities of best and nearby trajectories

We validate the least action approximation by systematically comparing the computed action against the numerical solution of the Fokker-Planck density. In this limit, the probability density *ρ*(*ϵ*_*f*_) ≡ *ρ*(*t*_*f*_, *ϵ*_*f*_) is expected to be proportional to exp[−*S*_*min*_(*ϵ*_*i*_, *ϵ*_*f*_)]. In Figure 4e, we plot the negative minimized action against the logarithm of the density for various final states. The strict linear relationship confirms that the system operates in a regime where the path integral is indeed dominated by the single most likely evolutionary path.

The approximation of Eq. (12) does not consider the contribution to the total probability of nearby trajectories, which are less likely than the least-action one, but more numerous. To estimate the entropic correction to the action computation due to this bundle of trajectories, we start by linearly expanding the log-argument of the integral—the action *S*[***ϵ***]—around the least-action trajectory 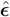 connecting *ϵ*_*i*_ at time *t* = 0 to *ϵ*_*f*_ at time *t* = *t*_*f*_. We define the Hessian matrix of the action through the second-order functional derivatives around this optimal trajectory:

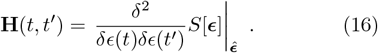

This Hessian matrix describes the local curvature of the trajectory landscape around the optimal path. Integrating over the fluctuations of ***ϵ*** leads to a Gaussian integral, resulting in an additive contribution to the least action,

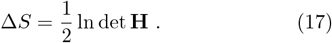

Physically, this correction is equal to (minus) the entropy of the bundle of trajectories surrounding the optimal path. A small determinant corresponds to a shallow valley in the action landscape (high entropy), allowing for significant fluctuations, whereas a large determinant indicates a steep valley where trajectories are tightly confined. Details about the calculation of the Hessian matrix and its determinant are found in Appendix D.

We validate this higher-order approximation in Figure 4f. In particular, we find that the small discrepancies observed in the zeroth-order approximation is perfectly explained by the entropic correction.

## IV. OPTIMAL PROTOCOLS

### A. Optimizing growth of high affinity trajectories

We seek the optimal antigen administration protocol *C*(*t*) that maximizes the likelihood of a specific evolutionary trajectory. In practice, we minimize the least action 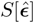 with respect to the control parameter *c*(*t*) = log *C*(*t*). We obtain two distinct regimes:

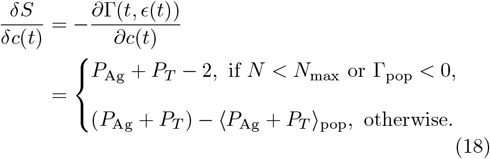

Here, Γ_pop_ ≡ ⟨log *P*_Ag_ +log *P*_*T*_⟩ _pop_ + *λ* represents the net population growth rate. Consequently, minimizing the action with respect to antigen concentration is equivalent to maximizing the instantaneous growth rate along the evolutionary trajectory. We define the *critical concentration, C*_0_, as the threshold where the net selection pressure exactly counterbalances the baseline rates of growth, mutation, and differentiation:

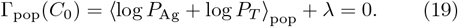

Below this threshold (*C < C*_0_), the system operates in an unconstrained growth regime where the population pressure term Ω(*C*) vanishes (Figure 5a). Since the Ag-binding and T-cell selection probabilities are bounded by unity, the derivative of the action remains negative (*P*_Ag_ + *P*_*T*_ − 2 *<* 0). Thus, the effective growth rate Γ increases monotonically with concentration up to *C*_0_, implying that the optimal protocol must maintain *C*(*t*) ≥ *C*_0_ to keep the population at carrying capacity *N*_max_ (Figure 5b). Conversely, above *C*_0_, the pressure term Ω(*C*) activates to enforce the population limit. In this regime, low-affinity B cells continue to benefit from higher antigen availability, but high-affinity clones experience a decrease in growth rate as the global population penalty Ω outweighs their saturated binding probabilities.

**FIG. 5.**
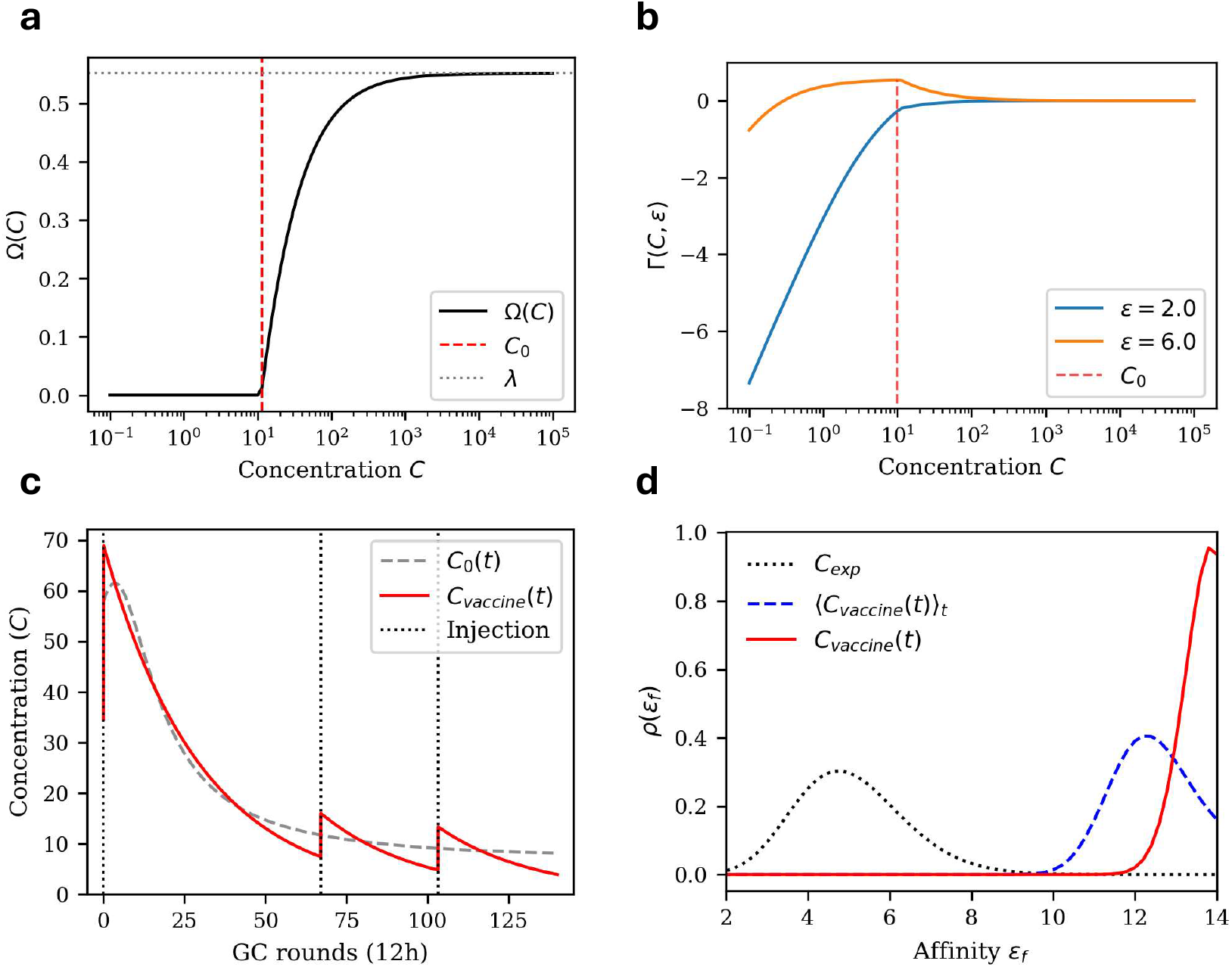
Optimal vaccination protocols. **a)** Carrying capacity pressure Ω(*C*) as a function of antigen concentration. The pressure term activates above a threshold *C*_0_, mediating the random removal of cells independently of affinity, and converges to the baseline growth rate *λ* in the limit of infinite concentration. **b)** Antigen concentration that maximizes the B-cell growth rate as a function of affinity. The optimum depends on the sign of the selection differential (*P*_Ag_ + *P*_*T*_) −⟨ *P*_Ag_ + *P*_*T*_⟩_pop_. Low-affinity B cells always benefit from higher antigen concentrations, while high-affinity cells achieve their maximal growth rate at an intermediate concentration *C*_0_. **c)** Optimal concentration protocol maximizing the probability of a trajectory terminating at *ϵ*_*f*_ = 6, alongside its best approximation using three tunable vaccine injections. **d)** Final B-cell affinity distributions comparing three protocols: the experimental concentration fitted to DeWitt et al., the theoretically derived optimal concentration, and a constant concentration set to the time-averaged value of the optimal protocol.

Since *C*_0_ depends on ⟨log *P*_Ag_ + log *P*_*T*_ ⟩_pop_, it varies as a function of time: when population affinity gets higher with GC rounds, the optimal concentration *C*_0_(*t*) decreases (Figure 5c).

### B. Optimal immunization scheme with discrete injections

The case of continuous antigen control is instructive but lacks realism. In reality, antigen is usually administered in a small number of injections and decays exponentially over time. For an immunization scheme consisting of *N*_inj_ injections, we can consider the antigen concentration to evolve as:

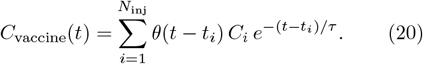

where *τ* = 30 GC cycles is the Ag decay time. For every injection *i* = 1, …, *N*_inj_, *t*_*i*_ corresponds to the injection time and *C*_*i*_ is the amount of antigen administered (related to the logarithmic amount by 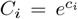). The function *θ*(Δ*t*) is the Heaviside step function.

To find the optimal injection schedule, we minimize the deviation between the discrete parametrization *C*_vaccine_(*t*) and the theoretically optimal continuous anti-gen control *C*_0_(*t*) derived in the previous section. We employ a gradient descent algorithm to minimize the Mean Squared Error (MSE) loss function defined over the simulation duration:

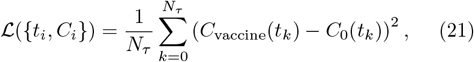

where the sum runs over all discrete time steps *t*_*k*_ of the simulation.

The results of this optimization are presented in Figure 5c. We observe that a simplified protocol consisting of *N*_inj_ = 3 tunable injections is sufficient to reproduce the key temporal features of the continuous least-action solution. The optimizer converges to a strategy that maintains antigen levels close to the critical selection threshold *C*_0_ during the intermediate rounds, preventing premature population extinction while sustaining selection pressure.

Finally, we validate the efficacy of this protocol by simulating the Fokker Planck equation under different immunization regimes. Figure 5d displays the final B-cell affinity distributions for the optimized vaccine schedule compared to the experimental baseline (fitted to data from DeWitt et al.[15]) and a constant concentration control fixed at the time-averaged value of the optimal input. The optimized discrete protocol induces a significant shift in the population distribution toward higher affinity states compared to the experimental trajectory. Notably, the time-varying protocol also outperforms the constant concentration baseline, demonstrating that temporal modulation of antigen availability—rather than total dosage alone—is critical for maximizing evolutionary gain.

## V. DISCUSSION

In this work, we have combined stochastic population dynamics with analytical optimization to define immunization protocols that maximize evolutionary gain. While previous studies [8–11] have established that antigen dosage is a critical determinant of affinity maturation outcomes, a rigorous derivation of the optimal temporal immunization profile has remained elusive. Our results provide a theoretical foundation for these observations, demonstrating that the most effective protocols rely on a dynamic modulation of selection pressure that guides the population along a “least-action” evolutionary trajectory. The main innovation of our approach is the reduction of complex agent-based dynamics to a tractable path-integral formalism. By mapping the stochastic evolution of B-cells to a Fokker-Planck equation, we analytically identified the “least-action” trajectory that maximizes the probability of reaching a final high-affinity state. We validated this framework by directly incorporating empirical measurements into our model parameters, specifically the activation energy and mutational effects derived from Deep Mutational Scanning (DMS) experiments by DeWitt et al. [15]. The remarkable agreement between our analytical predictions and the ensemble average of successful stochastic lineages confirms that, despite the inherent randomness of mutation, the mean evolutionary path is deterministic and predictable under strong selection.

Our findings corroborate and quantify the longstanding observation that affinity maturation is not monotonically improved by higher antigen dosages. Consistent with prior literature [2, 7, 8, 10, 11], we observe that excessive antigen concentrations relax selection pressure, allowing low-affinity clones to persist and dominate the population through sheer numbers rather than fitness. Conversely, insufficient antigen leads to overly stringent bottlenecks that extinguish the population before beneficial mutations can fixate. Our model formalizes this trade-off by identifying a critical, time-dependent concentration that keeps the system effectively at the “tipping point” of extinction. Beyond schedule timing, the availability of time-resolved sequence data now allows for the reconstruction of B-cell phylogenetic trees within individual subjects [16–19]. Tracking individual phylogeny could further improve our understanding of how vaccination protocols impact maturation by comparing these real histories to our theoretical “least-action” paths [20]. This comparison would help identify the specific selection pressures required to drive lineages toward high-affinity states, bridging the gap between clinical observation and deterministic evolutionary theory.

A key biological insight is that the optimal injection schedule maintains high selectivity over time. While our model does not incorporate all complexities of the immune response—such as the distinct recruitment kinetics of memory B cells [10, 21, 22]—it positions the injection protocol as a key factor, here supporting delayed second and third injections that effectively maintain a non-zero antigen concentration. This approach aligns with “slow delivery” immunization strategies, which have been shown to enhance germinal center responses in the context of HIV [23]. This principle is further supported by extensive research into SARS-CoV-2 strategies, where delayed dosing has consistently improved outcomes. In particular, extending the interval between doses has been associated with higher neutralizing antibody titers and stronger long-term protection against infection compared to standard intervals [24, 25].

To achieve mathematical tractability, we introduced necessary simplifications. We assumed a uniform proliferation rate, whereas biological evidence suggests that higher affinity B-cells may dwell longer in the Dark Zone and undergo more divisions [26, 27]. In our model, this fitness advantage is captured qualitatively through the selection probabilities *P*_Ag_ and *P*_*T*_. Furthermore, we modeled mutational effects using site-specific normal distributions fitted to DMS data. While we treat the distribution of mutational effects as independent of the current affinity state (similar to Wang et al. [12]), in reality, beneficial mutations likely become scarcer as the sequence approaches a fitness peak—a phenomenon known as diminishing returns [28]. Moreover, the modelization can be enriched with Ag depletion dynamics and the Germinal Center initiation process [10]. Finally, while we define the evolutionary target as high affinity to a single specific antigen, effective intervention against highly mutable pathogens like HIV [12–14] and SARS-CoV-2 [17] requires targeting breadth—the capacity to bind a diverse spectrum of viral variants— as recently modeled by Mahdisoltani et al. [29], who applied a path-integral framework to identify optimal vaccination protocols for the induction of broadly neutralizing antibodies.

## Author contributions

M.H., M.M., S.C. and R.M. designed the research, analyzed the results and wrote the paper. M.H. and M.M carried out numerical analysis and simulations.

## Appendix A Drift and diffusion coefficients

If we exclude lethal mutations (their effect is already accounted for in the growth term *λ*) then the probability for a cell to undergo a binding-energy change *ϵ* → *ϵ* + Δ*ϵ* in one round is given by the kernel *K*:

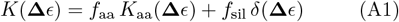

Here, 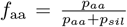 and *f*_sil_ = 1 − *f*_aa_ are the frac-tions of affinity-affecting and silent mutations, respectively. The kernel *K*_aa_ is a normal distribution with mean *µ*_**M**_ and variance 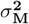 while *δ* is the Dirac delta function centered at zero.

In the limit of large population size, affinity-affecting mutations can be described by a diffusion process. The effective drift and diffusion coefficients are then:

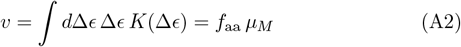

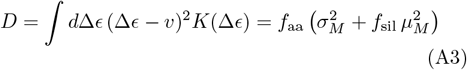

For the numerical simulation of the Fokker-Planck equation, we employed a time step of *δt* = 0.001 and a spatial discretization of *δϵ* = 0.05.

## Appendix B Path–integral formulation

For a short time increment *δt* ≪ 1, we formally write the evolution of the density as:

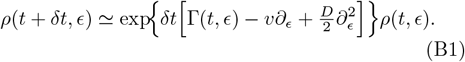

We insert the Dirac delta in Fourier representation,

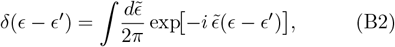

and evaluate (B1) at an initial position *ϵ*_*i*_ and a final position *ϵ*_*f*_ ≡ *ϵ*_*i*_ + Δ*ϵ*. We obtain:

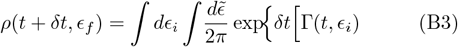

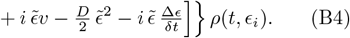

Equation (B4) is the *one–step propagator* between two consecutive time slices of duration *δt*.

We partition the total interval [*t*_0_, *t*_*f*_] into *T* = *t*_*f*_ − *t*_0_ = *Nδt* steps labelled by an integer *τ* = 0, …, *N*_*τ*_ − 1. Repeatedly inserting (B4) yields the joint integral over all intermediate positions {*ϵ*_*τ*_} and their conjugate Fourier modes 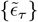:

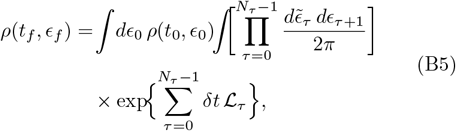

where we introduced the *discrete Lagrangian*:

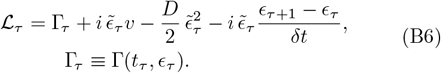

Equation (B5) is *quadratic* in 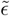, which can be integrated out analytically using the Gaussian identity:

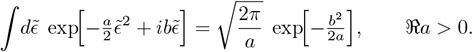

Setting *a* = *Dδt* and *b* = *δt v* − (*ϵ*_*τ*+1_ − *ϵ*_*τ*_), and performing all *N* integrations gives the purely *real* sum:

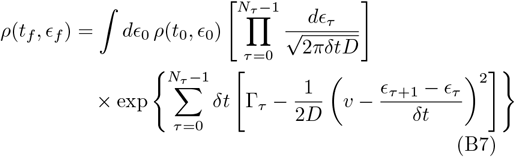

Taking the continuum limit *δt* → 0 (*N*_*τ*_ at fixed *T* → ∞), we identify *ϵ*_*τ*_ → *ϵ*(*t*) and the exponent becomes a time integral. We thus obtain the path–integral representation:

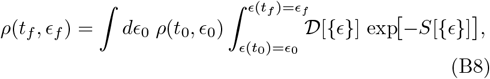

with the real action:

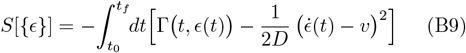

For the numerical computation of action, we considered *N*_*τ*_ = 1000 intermediate positions.

## Appendix C Least action

The dominant contribution to (B8) in the weak–noise limit arises from the trajectory that minimizes *S*[*ϵ*].

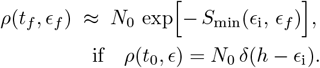

We identify the integrand of the action *S*[*ϵ*] = *dt* as the effective Lagrangian:

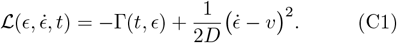

Extremizing the action yields the Euler–Lagrange equation:

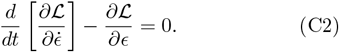

Computing the partial derivatives,

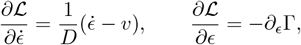

and substituting them back gives:

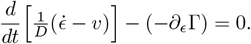

This simplifies to the second-order ODE:

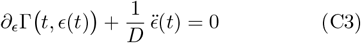

In our numerical experiment, the trajectory must satisfy boundary conditions:

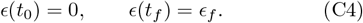

It can be solved with standard BVP solvers (e. g. `scipy.integrate.solve bvp`).

## Appendix D Entropic corrections

We evaluate the probability density *ρ*(*T, ϵ*_*T*_), when starting from the wildtype *ϵ*_*i*_ = 0, by integrating over all possible trajectories ***ϵ***. While the path integral is dominated by the least-action trajectory 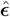, fluctuations around this optimal path contribute to the total probability (the entropic contribution).

We decompose a general trajectory ***ϵ*** into the optimal path 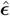 and a fluctuation vector Δ***ϵ***:

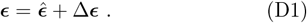

We expand the action *S*[***ϵ***] to the second order around 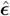 using a functional Taylor expansion:

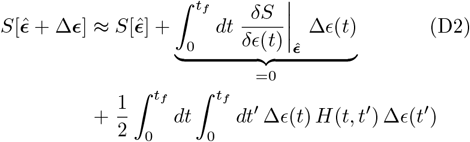

The first-order term vanishes because 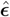 is a stationary point of the action (satisfying the Euler-Lagrange equations). The second-order term is governed by the Hessian matrix **H**, which describes the local curvature of the fitness landscape.

The elements of the Hessian **H** matrix, see Eqn. (16), can be expressed in a discrete approximation with step *δt*, with *N*_*T*_ = *t*_*f*_ */δt* time steps between the initial and final times. This yields a (*N*_*T*_ × *N*_*T*_)–symmetric tridiagonal matrix with components:

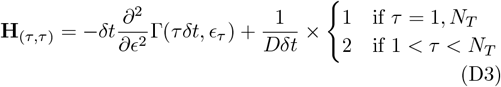

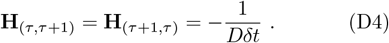

To have a well defined limit when *δt* → 0,it is appropriate to rescale the Hessian matrix as **H** → *D* · *δt* · **H**.

We therefore obtain:

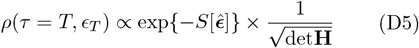

up to an irrelevant multiplicative factor.

In practice, this involves computing the determinant of a Hessian of size *N*_*T*_ × *N*_*T*_, where we choose *N*_*T*_ = 1000. While we utilized the general-purpose `numpy.linalg.slogdet` solver for our calculations, the tridiagonal structure of the Hessian allows for significantly more efficient computation [30].

## Appendix E Concentration optimisation

We control the time–dependent antigen concentration through its logarithm

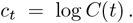

The action minimised by the least-action trajectory *ϵ*(*t*) is

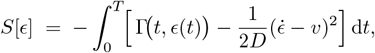

with growth term

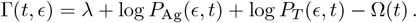

Because *c*_*t*_ enters only through Γ,

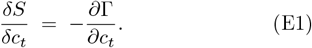

Differentiating with respect to *c*_*t*_,

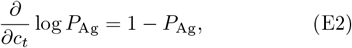

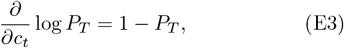

so that

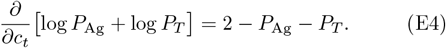

The size constraint is enforced by

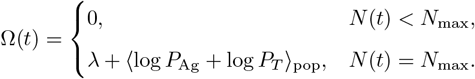

Hence

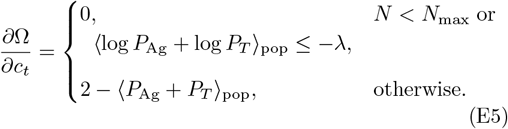

Combining (E1), (E4) and (E5),

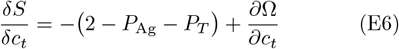

which becomes, explicitly,

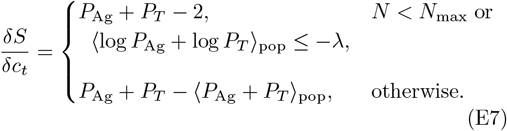

